# Deciphering human ribonucleoprotein regulatory networks

**DOI:** 10.1101/295097

**Authors:** Neelanjan Mukherjee, Hans-Hermann Wessels, Svetlana Lebedeva, Marcin Sajek, Mahsa Ghanbari, Aitor Garzia, Alina Munteanu, Thalia Farazi, Jessica I Hoell, Kemal Akat, Thomas Tuschl, Uwe Ohler

**Affiliations:** Department of Biochemistry and Molecular Genetics, RNA Bioscience Initiative, University of Colorado School of Medicine, Aurora, Colorado 80045, USA.; Berlin Institute for Medical Systems Biology, Max Delbrück Center for Molecular Medicine, Berlin, Germany.; lnstitute of Human Genetics, Polish Academy of Sciences, Poznan, Poland; Howard Hughes Medical Institute and Laboratory for RNA Molecular Biology, The Rockefeller University, 1230 York Ave, Box 186, New York, NY 10065; Department of Pediatric Oncology, Hematology and Clinical Immunology, Center for Child and Adolescent Health, Medical Faculty, Heinrich Heine University of Dusseldorf, Dusseldorf, Germany

**Author notes:** Current address: Juno Therapeutics, 400 Dexter Avenue North, Seattle, WA 98109.

## Abstract

RNA-binding proteins (RBPs) control and coordinate each stage in the life cycle of RNAs. Although *in vivo* binding sites of RBPs can now be determined genome-wide, most studies typically focused on individual RBPs. Here, we examined a large compendium of 114 high-quality transcriptome-wide *in vivo* RBP-RNA cross-linking interaction datasets generated by the same protocol in the same cell line and representing 64 distinct RBPs. Comparative analysis of categories of target RNA binding preference, sequence preference, and transcript region specificity was performed, and identified potential posttranscriptional regulatory modules, i.e. specific combinations of RBPs that bind to specific sets of RNAs and targeted regions. These regulatory modules encoded functionally related proteins and exhibited distinct differences in RNA metabolism, expression variance, as well as subcellular localization. This integrative investigation of experimental RBP-RNA interaction evidence and RBP regulatory function in a human cell line will be a valuable resource for understanding the complexity of post-transcriptional regulation.

## Introduction

Of the 20,345 annotated protein-coding genes in human, at least 1,542 are RNA-binding proteins (RBPs) (Gerstberger et al., 2014). RBPs interact with RNA regulatory elements within RNA targets to control splicing, nuclear export, localization, stability, and translation (Moore, 2005). RBPs have specificity to bind one or multiple RNA categories, including messenger RNA (mRNA) and diverse categories of non-coding RNA such as ribosomal RNA (rRNA), transfer RNA (tRNA), small nuclear and nucleolar RNA (snRNA/snoRNA), microRNA (miRNA), and long non-coding RNA (lncRNA). Mutations in RBPs or RNA regulatory elements can result in defects in RNA metabolism that cause human disease (Cooper et al., 2009; Fredericks et al., 2015).

A standard technique for *in vivo* global identification of RBP-RNA interaction sites consists of immunoprecipitating the ribonucleoprotein (RNP) complex, isolating the bound RNA, and quantifying the RNA targets by microarrays or deep sequencing (Tenenbaum et al., 2000; Zhao et al., 2010). The introduction of cross-linking prior to immunoprecipitation (CLIP) as well as RNase digestion enabled the biochemical mapping of individual interaction sites (Ule et al., 2003). Subsequent modifications to CLIP increased the resolution of the interaction sites (Hafner et al., 2010; König et al., 2010). One of these methods, photoactivatable ribonucleoside-enhanced cross-linking and immunoprecipitation (PAR-CLIP), utilizes 4-thiouridine or 6-thioguanosine combined with 365 nm UV crosslinking to produce single-nucleotide RBP-RNA interaction evidence that is utilized to define binding sites (Corcoran et al., 2011; Garzia et al., 2017b; Hafner et al., 2010).

Experimentally-derived RBP binding sites provide valuable functional insights. First, they can reveal the rules for regulatory site recognition by the RBP, whether due to sequence and/or structural characteristics. Second, the region and position of the interaction sites of an RBP within transcripts provides insights into its role in RNA metabolism and its subcellular localization. For example, if most of the mapped interaction sites are intronic and adjacent to splice sites, the RBP is highly likely to be a nuclear splicing factor rather than a cytoplasmic translation factor. Finally, these data reveal the target transcripts and therefore the potential biological role for the RBP.

Throughout the life of an RNA, interactions with many different RBPs determine the ultimate fate of the transcript. Even though profiling of the interaction sites of a single RBP is clearly powerful, it does not provide information on other RBPs potentially targeting the same RNA or on other regulatory elements within the RNA. Small comparative efforts focusing on the regulation of splicing, 3’ end processing, RNA stability by AU-rich elements, and miRNA-mediated silencing have demonstrated the value of integrating interaction sites from multiple RBPs (Martin et al., 2012; Mukherjee et al., 2014; Pandit et al., 2013; Zhang et al., 2010).

Therefore, a large-scale comparative examination of interaction sites for many RBPs will yield valuable knowledge regarding the architecture and determinants of RNA regulatory networks.

At least 173 PAR-CLIP experiments have been performed in HEK293 cells to date, laying the groundwork for a large-scale integrative analysis and complementing efforts of ENCODE, which focused on other cell types and utilized other CLIP protocols (Van Nostrand et al., 2016). We describe a concerted effort to identify and uniformly process all high-quality PAR-CLIP data sets by evaluating the characteristic T-to-C transitions induced by photocrosslinking. Using the resulting compendium of high-quality in vivo RBP interaction maps from the same cell line enabled us to determine the relationship between RBPs with respect to their preferred category of target RNA and any underlying sequence specificity. We uncovered regulatory modules reflected by combinatorial binding events, and assessed their role and functional implications on RNA metabolism. Finally, our results support the role of RBPs in buffering gene expression variance.

## Results

### A high-quality map of in vivo RBP-RNA interactions across 64 proteins

In order to generate a comprehensive quantitative resource of RBP-RNA interactions within a human cell line, we identified 166 published PAR-CLIP data sets performed predominantly in HEK293 cells, and added 7 new libraries generated in our laboratories (Sup Table 1). Typically, these datasets were generated using transgenic HEK293 cell lines in which each individual RBP was FLAG-tagged and recombined into the same chromosomal locus containing a strong promoter. In this way, the expression of each RBP as well as the strength of its immunoprecipitation were generally comparable. Furthermore, the availability of orthogonal transcriptome-wide datasets quantifying individual steps of RNA metabolism made HEK293 cells ideal for examining the functional characteristics of RNA targets (Mukherjee et al., 2017).

Each of the 173 PAR-CLIP libraries generated in HEK293 were subject to a stringent analysis strategy to retain high-quality datasets (Supplemental Table 1). First, each library was analyzed using the PAR-CLIP Suite v1.0 (https://rnaworld.rockefeller.edu/PARCLIP_suite) (Garzia et al., 2017b) to discriminate significant target RNA categories from non-crosslinked background RNA categories populated by fragments of abundant cellular RNAs (see Methods, Supplemental Fig. 1A). Next, we defined binding sites based on the local density of T-to-C transitions using PARpipe (https://github.com/ohlerlab/PARpipe) (Corcoran et al., 2011) and only retained those libraries with sufficiently high read counts and T-to-C transition specificity compared to a deeply sequenced background reference library (Supplemental Fig 1b) (Friedersdorf and Keene, 2014). Since the immunoprecipitation step was omitted in this reference library it served as an effective comparison point to score read count and T-to-C transition for all RBPs. Finally, for RBPs with more than 3 libraries available, outlier libraries exhibiting poor correlation of 6-mer frequencies were excluded (Supplemental Fig 1d, e). This resulted in 114 libraries corresponding to 64 RBPs that were the basis for downstream analysis. There were eight RBP families represented by two or more RBPs.

### Grouping RBPs by annotation category and positional binding site preferences

As first step to describe RBP-RNA regulatory networks, we determined the relative binding preference of each RBP for specific target RNA annotation categories (Supplemental Table 2). For each library, we calculated an RNA annotation category preference value, defined as the difference in the fraction of T-to-C reads per annotation category between each RBP library and the reference library. We performed hierarchical clustering of RBPs by annotation category preference, using Ward’s method and Euclidean distances. This yielded eight clusters of binding preference (Figure 1a – orange line demarcates cluster definitions) with varying enrichment or depletion for individual or combinations of specific annotation categories. For each of these clusters, we compiled a detailed table summarizing the reported functions for each of the RBPs (Table 1). Taken together, clustering by RNA annotation category separated RBPs into groups according to their known subcellular localization and functions.

**Figure 1:**
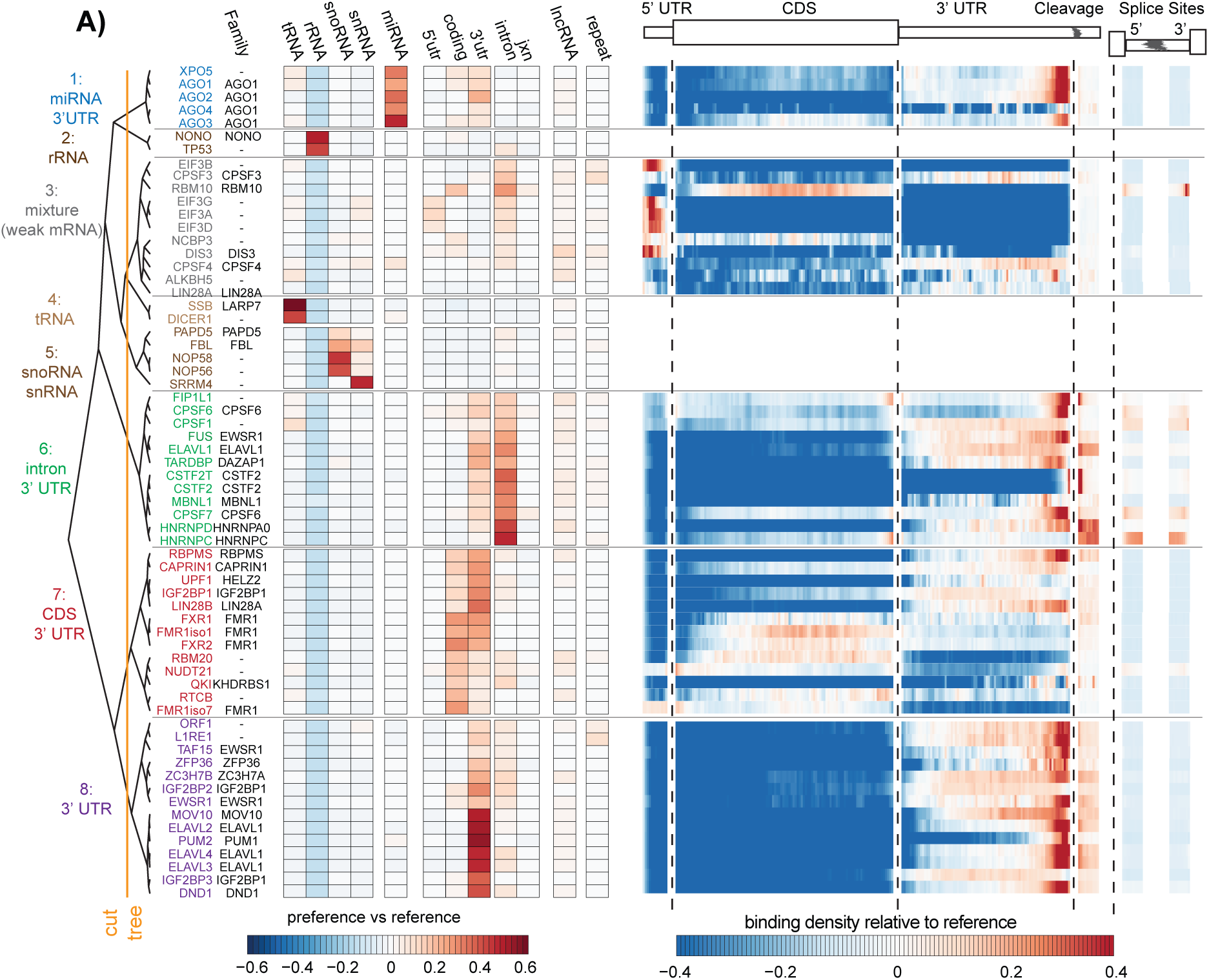
RBP analyzed and binding preferences by RNA category. A) Heatmap of reference normalized annotation category preference for each RBP clustered into 8 branches by color (left). The heatmap represents the difference in the proportion of sites for a given annotation category in the RBP library versus the reference library. Heatmap of the reference library normalized relative positional binding preference of the 55 RBPs with enriched binding in at least one mRNA-relevant annotation category per branch (right). RBP-specific binding preferences were averaged across selected transcripts (see methods). The relative spatial proportion of 5’UTR, coding regions and 3’UTR were averaged across all selected transcript isoforms. For TES (regions beyond transcription end site), 5’ splice site, and 3’ splice site, we chose fixed windows (250nt for TES and 500nt for splice sites). For each RBP, meta-coverage was scaled between 5’UTR to TES. The 5’ and 3’ intronic splice site coverage was scaled separately from other regions but relative to each other.

Three of the eight clusters (clusters 2, 4, and 5) contained nine RBPs that exhibited preference for categories of non-coding RNA (rRNA, snRNA, snoRNA, and tRNA), but not mRNA, precursor mRNA (pre-mRNA), or lncRNA. The remaining five clusters contained 55 RBPs exhibiting preference for binding to mRNA, pre-mRNA and long-noncoding RNA (lncRNA) annotation categories. The RBPs in clusters 1, 6, 7, and 8 exhibited strong preferences for various mRNA annotation categories. The RBPs in cluster 3 did not exhibiting strong preference for specific mRNA annotation categories. Additionally, for each of the RBPs in the cluster, we performed a positional meta-analysis of binding sites with respect to major transcript landmarks within target mRNAs. Many of the RBPs also showed strong preferences for binding to specific positions within mRNAs relating to their role in specific steps of mRNA processing (Table 1).

We hypothesized that target annotation category preferences and positional binding preferences should reflect subcellular localization of the RBP and its role(s) in mRNA processing. Cluster 6 contained twelve RBPs and exhibited strong preference for intronic regions and to a lesser degree 3’ UTRs of mRNAs and lncRNAs. The intronic preference was consistent with the predominantly nuclear localization of these RBPs and the pre-mRNA splicing process. ELAVL1, which is the sole member of the ELAVL1 family of RBPs that is predominantly localized in the nucleus but capable of shuttling to the cytoplasm, exhibited positional binding flanking the end of the 3’ UTR and for 5’ and 3’ splice sites. Cluster 8 contained fourteen RBPs and exhibited distinct preference for 3’ UTR regions. This included the unpublished and predominantly cytoplasmic ELAVL1 family members, ELAVL2, ELAVL3, and ELAVL4, which exhibited a strong positional preference for binding in the distal region of the 3’ UTR and acting predominantly on mature mRNA (Mansfield and Keene, 2012). In summary, the annotation category preferences and positional binding preferences implicated the specific steps of mRNA processing the RBPs potentially regulate.

### The spectrum of RNA sequence specificity

RBPs exist on a spectrum of specificity depending on a variety of primary and secondary structure features (Jankowsky and Harris, 2015). Here, our goal was to identify the RBPs with substantial primary sequence specificity and then examine their sequence preference. For each of the 55 RBPs, we counted all possible 6-mers using Jellyfish (Marçais and Kingsford, 2011) for the reads contributing to PARalyzer-defined binding sites. We observed 6-mer frequencies ranging as high as 512-fold to as low as 5-fold over a uniform distribution of 6-mers (Supplemental figure 2a). In contrast, our reference background library exhibited 16-fold enrichment of at least one 6-mer compared to uniform. AGO1-4 libraries were excluded from 6-mer analysis due to the overwhelming sequence contribution from crosslinked miRNAs. Twenty-seven RBPs did not have a single 6-mer found at higher frequency than present in the reference sample. Amongst these RBPs established or expected to display low sequence-specificity were the RNA helicase MOV10, the nuclear exosome component DIS3, and the EIF3 complex translation initiation factors.

For each of the 24 RBPs with stronger sequence enrichment than the reference library, we clustered the top 5 sequences enriched over the reference library (Figure 2). Our results recapitulated the sequence preference for the RBPs in this group with well-characterized sequence motifs (detailed in Table 2). The ELAVL1 family proteins, which bound to different regions and positions of mRNA, showed similar preference for U- and AU-rich 6-mers, while ZFP36 only enriched a subset of the AU-rich 6-mers (Mukherjee et al., 2014). Complementing the 6-mer enrichment analysis, we performed motif analysis for each RBP library with the motif finding algorithm SSMART (sequence-structure motif identification for RNA-binding proteins, (Munteanu et al., 2018)) (Supplemental Fig 2b). For most RBPs, we observed strong concordance between the two analyses. RBM20 was a clear exception, for which we observed the established UCUU-containing motifs (Maatz et al., 2014) with SSMART, but a GA-rich sequence in the 6-mer enrichment analysis. However, we do observe UCUU-containing motifs in the top 15, but not top5 6-mers for RBM20. Altogether, our analysis was remarkably consistent with previously reported motifs in spite of differences in data processing and analysis (detailed Table 2).

**Figure 2:**
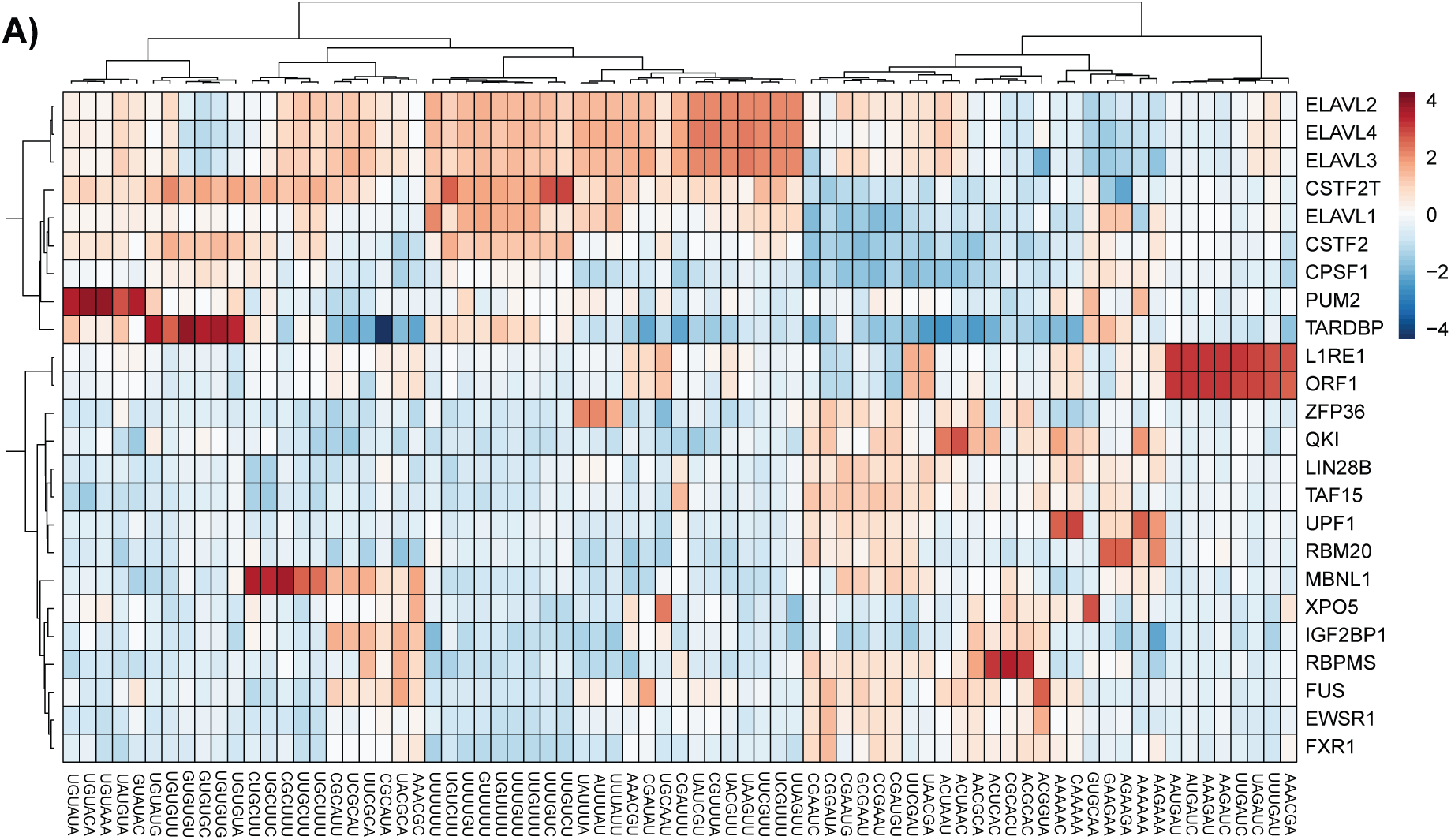
RBP binding sequence specificity and elements. A) Heatmap of reference normalized 6-mer enrichment for top 5 enriched 6-mers for each RBP in the set of RBPs exhibiting more sequence specificity than the reference.

### Identification of RNA regulatory modules

To understand the functional impact of co-regulation by multiple RBPs, we analyzed the co-variation in binding patterns of all 55 RBPs across 13,299 target RNA encoding genes to probe for the existence of regulatory modules, i.e., specific subsets of RNAs implicated in similar function bound by subsets of RBPs. To this end, we employed Factor Analysis (FA), which reduces a large number of observed variables to a smaller number of latent *factors*. Here, our observed variables represented the normalized RBP binding (see methods) for each of the 55 RBPs across all target RNA encoding genes (n=13,299). The latent *factors* represented similar binding patterns to RNA targets by one or more of the 55 RBPs. RBPs exhibiting high loadings for the same *factor* would have very similar binding patterns to RNA targets. Importantly in this framework, a single RBP could be assigned to multiple *factors*, just as a single RBP can participate in multiple RNPs and regulate different aspects of RNA metabolism.

The FA model decomposed the 55 x 13,299 normalized RBP binding matrix into a 55 x 10 factor loading matrix (representing the strength of the dependence of each of the 55 RBP target RNA binding pattern on each of the 10 *factors*), a 13,299 x 10 factor score coefficient matrix (representing the dependence between the binding of the 13,299 target RNA encoding gene and each of the 10 *factors*), and residual error (Supplemental Fig 3a and methods). Cumulatively, the FA model explained ~60% of the variance in the observed data. The remaining unexplained variance was expected due to the challenges of integrating data sets of varying depth and quality, in spite of our efforts to control these aspects. The communality, which is the amount of variance explained by the model for each RBP-binding variable, varied drastically for all 55 RBPs; the model explained at least 80% of the variance in enrichment scores for 12 RBPs, and at least 50% of the variance in enrichment scores for 30 RBPs (Supplemental Figure 3b). RBPs with lower communality often coincided with shallow depth of their PAR-CLIP libraries.

The FA model also uncovered interesting parallels between the similarity in the binding of target RNA encoding genes and the target annotation category preferences (from Figure 1a). We observed that individual *factors* contained RBPs that preferred binding to either mature (Factors 1, 3, 4, 5, 8) or precursor transcripts (Factors 2, 6), reflecting involvement in different stages of RNA metabolism (Figure 3a). Furthermore, individual *factors* contained RBPs exhibiting similar patterns of binding to specific regions of the mRNA (i.e., intron, coding, 3’ UTR). Indeed, RBPs from the same family, or known to regulate a specific aspect of RNA processing, had high loadings for the same *factors*. For example, the ELAVL1 family members were associated with Factor 1; the AGO1 family were associated with Factor 3; the IGF2BP1 family were associated with Factor 4; the FMR1 family had were associated with Factor 5 and Factor 8; LINE-1 encoded proteins were associated with Factor 7. One of the unanticipated associations was that of HNRNPC with Factor 2, which contained mainly cleavage and polyadenylation factors. Interestingly, HNRNPC was shown to interact with U-rich sequences downstream of a viral poly-adenylation signal nearly three decades ago (Wilusz et al., 1988), and more recently, to repress cleavage and poly-adenylation in humans (Gruber et al., 2016). These examples highlight the specific testable hypotheses generated by an integrative analysis that are not necessarily obvious when examining a single RBP in isolation.

**Figure 3:**
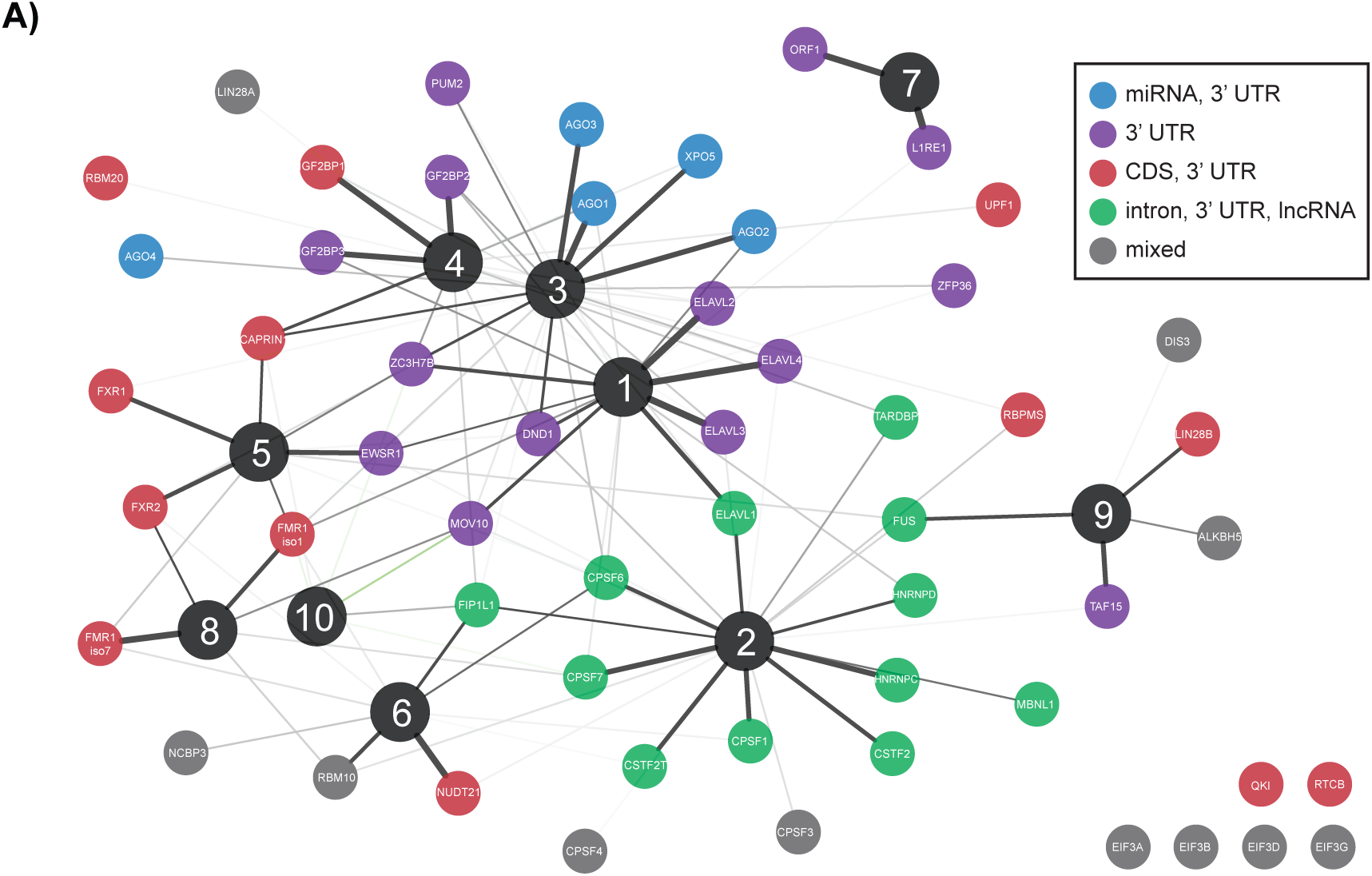
RNA regulatory modules. A) Factor analysis of target RNA encoding genes binding normalized by the reference library and expression for the 55 RBPs binding to mRNAs and lncRNAs for 13,299 genes (see ‘factor analysis’ section in methods for details). Spring-embedded graph of the factor loading matrix, indicating the association between each of the 55 RBPs and one of the 10 factors. Nodes color-coded by RNA annotation category preference cluster membership from figure 1. Edge width scales with factor loadings (thicker edge = higher factor loading = stronger association). Only edges with a factor loading > 0.2 (positive values in black) or < -.2 (negative values in green) depicted.

By clustering the factor score coefficients, i.e. the specific linear combination of RBP binding for that target RNA, we identified target RNA encoding genes constituting putative regulatory modules associated with a given *factor*. Therefore, each regulatory module was associated with an RBP component (the subset of RBPs exhibiting similar binding pattern) and a RNA component (the subsets of target RNA encoding genes bound by those RBPs). These regulatory modules did not imply physical interactions between RBPs; rather, it identified RBPs that may cooperate in controlling RNA metabolism for specific subsets of RNA targets, possibly across cellular compartments. Almost a quarter of the target RNA encoding genes (3,180/13,299) were assigned to regulatory modules by exhibiting high factor score coefficients for a single *factor* (Supplemental figure 3c). We did not identify target RNA encoding genes with high factor score coefficients for Factor 9 or 10. The remaining target RNA encoding genes did not exhibit high factor score coefficients for any specific *factor* in our analysis, suggesting that the targets were either not bound by specific combinations of these RBPs, bound broadly by all RBPs, or not bound by the subset of RBPs in the analysis. As such, we labeled this target RNA encoding gene category as “non-specific”. The RNA regulatory modules encoding genes were enriched for different GO categories. Factor 1 RNA regulatory modules were enriched for ‘AU-rich element binding’ and Factor 3 RNA regulatory modules were enriched for ‘gene silencing by miRNA’; AU-rich RBPs and AGO proteins were strongly associated with Factor 1 and Factor 3, respectively. This was consistent with the recurrent observation that RBPs target the mRNAs encoding themselves (Pullmann et al., 2007; Tenenbaum et al., 2000). In turn, the RNAs encoding “non-specific” genes contained ribosomal proteins and mitochondrial electron-transport proteins.

### RNA regulatory modules underlie distinct patterns of RNA metabolism

In order to test the functional relevance of these RNA regulatory modules, we reasoned that perturbation (change of protein abundance or activity) of an RBP will lead to pronounced effects only for the RNA regulatory modules assigned to the specific *factor(s)* that RBP is associated with. We examined mature and precursor RNA expression changes induced by siRNA knockdown of ELAVL1 (Kishore et al., 2011). ELAVL1 was strongly associated with both Factor 1 and Factor 2, which exhibited RNA targeting patterns for mature or precursor RNAs, respectively. Concordantly, Factor 1 associated RNA regulatory modules, but not Factor 2 RNA regulatory modules, exhibited ELAVL1-dependent stabilization of mature RNA (Figure 4a). Likewise, Factor 2 RNA regulatory modules exhibited a more pronounced ELAVL1-dependent stabilization of precursor RNA than Factor 1 RNA regulatory modules (Figure 4b). Each human ELAV1 family protein contains three RRM domains (>90% sequence identity), but the hinge region between the second and third RRM of ELAVL1 contains a shuttling sequence responsible for its nuclear localization (Fan and Steitz, 1998). Due to the lack of this shuttling sequence, ELAVL2/3/4 are predominantly cytoplasmic and were strongly associated with Factor 1, but not Factor 2. Taken together, the model was able to correctly identify and distinguish ELAVL1-dependent stabilization of both precursor and mature RNA (Lebedeva et al., 2011; Mukherjee et al., 2011).

**Figure 4:**
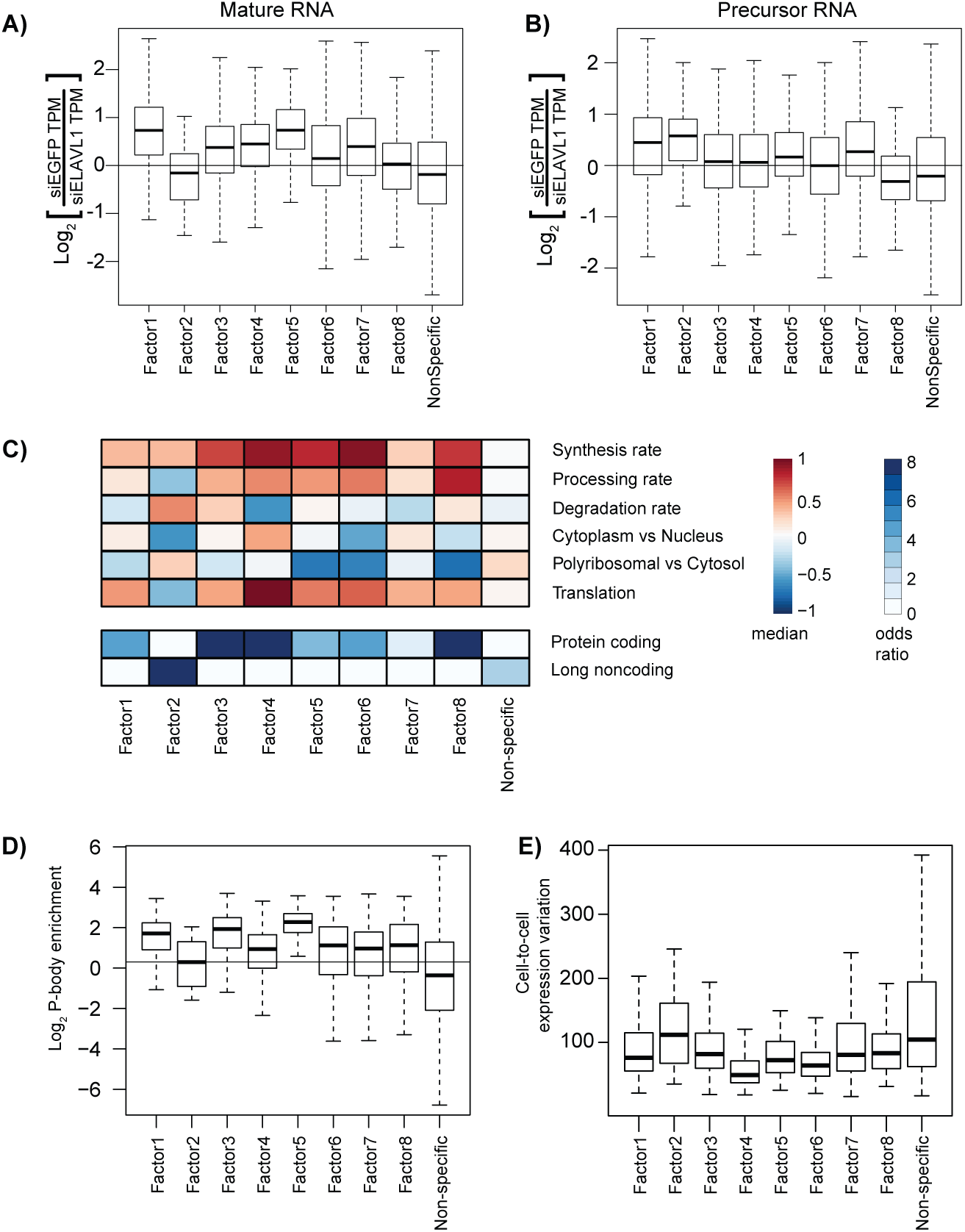
Functional characterization of RNA regulatory modules. A) The difference in either A) primary or B) mature RNA expression (transcripts per million) upon ELAVL1 knockdown by siRNA treatment (y-axis), specifically the log_2_[siRNA EGFP TPM]-log_2_[siRNA ELAVL1 TPM], foreach gene set. C) Heatmap of the median value of synthesis rate, processing rates, degradation rates, cytoplasmic versus nuclear localization, polyribosomal versus cytoplasmic localization, and translational status from ribosome profiling data for each gene set (top). Heatmap of the odds-ratio of the overlap between factor associated gene sets with annotation (bottom). D) Box- and-whisker plot for each gene set of the enrichment in P-bodies. E) Box-and-whisker plot for each gene set of the coefficient of variation across 25 individual HEK293 cells.

We hypothesized that the subsets of RNAs assigned to the different regulatory module would exhibit differences in RNA metabolism driven by the RBPs in the *factor* associated with the regulatory module. Therefore, we compared six aspects of RNA metabolism previously quantified in HEK293 cells (Mukherjee et al., 2017), for each of the RNA regulatory modules associated with each of the *factors*. The *factor*-associated RNA regulatory modules exhibited very distinct RNA metabolic profiles compared to each other and to non-specific category (Figure 4c, Supplemental Figure 4a). Factor 2 RNA regulatory modules, which was the only factor associated with RBPs binding to precursor mRNA and lncRNA, had low processing rates, high degradation rates and their encoded RNAs were preferentially localized in the nucleus versus the cytoplasm. Factor 2 RNA regulatory modules were strongly enriched for lncRNAs (Figure 4d). Indeed, these genes strongly overlapped with a set of lncRNAs likely to be functional (Supplemental figure 4b) (Mukherjee et al., 2017).

We also examined regulatory differences in RNA metabolism for genes associated with cytoplasm-enriched factors. For example, factor 1 RNA regulatory modules were more stable than Factor 3 RNA regulatory modules (Figure 4c). Factor 1 was strongly associated with ELAVL1 family proteins, which stabilize target mRNAs. Factor 3 was strongly associated with AGO1 family proteins, which execute miRNA-mediated degradation of target mRNAs. Additionally, Factor 4 RNA regulatory modules, which are bound by IGF2BP1 family proteins, were highly synthesized, processed, stabilized, and translated (Figure 4c). The RNA targets of IGF2BP1 family RBPs were strongly localized to the ER (Supplemental Figure 4c) (Jønson et al., 2007), which is also consistent with the proposed role of IGF2BP1 family proteins for RNA localization and translation (Farina et al., 2003; Nielsen et al., 2001). Although correlative, these results indicate that different RBP binding patterns beget different consequences for RNA metabolism.

Specific RNA regulatory modules also exhibited preferential localization to processing bodies (P-bodies), which are cytoplasmic granules associated with translational repression (Sheth and Parker, 2003). Namely, Factor 3 RNA regulatory modules, which were strongly associated with the AGO1 family, were the most strongly enriched for localizing to P-bodies according to a recent study characterizing the transcriptome and proteome of P-bodies, and the AGO2 protein itself was 90-fold enriched (Hubstenberger et al., 2017). Similarly, Factor 5 RNA regulatory modules, which were strongly associated with the FMR1 family, were also enriched for localizing in P-bodies, along with the FMR1 protein (16-fold enriched). In contrast, the non-specific category was depleted from P-bodies.

Fine-tuning of gene expression has been postulated to be an important function of post-transcriptional regulation by RBP and miRNAs. Therefore, we examined the cell-to-cell variability in gene expression across 25 individual HEK293 cells with respect to the RNA regulatory modules. The single-cell RNA-seq data was very deeply sequenced and generated using the massively parallel single-cell RNA-sequencing (MARS-Seq) protocol (Guillaumet-Adkins et al., 2017). Most RNA regulatory modules exhibited lower expression variability than the non-specific category (Figure 4e). In particular, Factor 4 RNA regulatory modules exhibited the lowest variation and highest median expression across the 25 cells (Supplemental Figure 4d). These results supported the broad notion that post-transcriptional gene regulation generally confers robustness and fine-tuning of gene expression.

## Conclusion

Our study presents a curation of existing datasets, followed by systematic analysis of high-quality and high-resolution RBP-RNA interaction data. We focused on the RBPs that preferentially bound to mRNA and lncRNA and examined their sequence specificity and sequence motif preferences. Our survey of the RBP regulatory landscape identified the most prevalent subsets of RNAs targeted by a specific subset of RBPs, which we refer to as RNA regulatory modules.

We utilized high quality PAR-CLIP datasets for which the immunoprecipitation was generally comparable due to fact most RBPs were FLAG-tagged. Nevertheless, several caveats associated with the interpretation of this analysis need to be pointed out. Despite several measures of quality control to decide which datasets to include in our analysis, the libraries varied greatly in depth, quality, digestion biases and potentially other confounding variables with respect to the protocol. The FA model quantitatively assessed the degree to which we could explain the full complement of RBP-RNA target binding patterns. These confounders undoubtedly contributed to the ~40% of variance not explained by the FA model. In comparison, the ENCODE eCLIP datasets (Van Nostrand et al., 2016) are likely to suffer from different confounders: they were generated using one consistent experimental protocol but used antibodies against endogenous proteins expressed at varying levels, and for which IP efficiency can vary greatly in spite of the quality control performed (Sundararaman et al., 2016). Essentially, this represents the trade-offs in experimental design between analyzing the endogenous protein compared to an epitope-tagged protein. Modifying the genomic loci of the protein to engineer an endogenous epitope tagged RBP is a very promising strategy.

Assuming the RBPs investigated here are a representative sample of the ~1,542 RBPs encoded in the human genome, there may be an astounding number of RBPs with substantial primary sequence specificity. However, the degree of sequence specificity is determined by the nature of the RBP-RNA interaction, which can be quite extensive and specific, as in the case of Pumilio, or minimal and non-sequence specific, as in the case of an RNA-helicase. An interesting exception were the A-rich sequences enriched by UPF1, which is an RNA helicase and therefore unlikely to exhibit strong sequence specificity. One possible explanation is that such sequences may represent pre-mature polyA tail recognition involved in aspects of ribosome quality control demonstrated in yeast (Koutmou et al., 2015) and human cells (Garzia et al., 2017a). Likewise, more examples of unanticipated sequence enrichments may shed light on novel RNA regulatory mechanisms.

Our FA model was able to identify distinct RBP-RNA target regulatory modules. At the very minimum, 25% of target RNA encoding genes were assigned to RNA regulatory modules. This is very likely an underestimation due to noisy data and a biased, far from complete sampling of RBPs. However, there is likely to be a subset of genes for which post-transcriptional gene regulation indeed plays a negligible role, at least in HEK293 cells. Furthermore, a small number of RBPs in our analysis are not endogenously expressed in HEK293 and their natural expression is tissue-specific and/or context-dependent. The approach presented here can scale to binding data for all ~700 RBPs experimentally shown to be associated with poly-adenylated RNA in HEK293 cells or even ~1,542 known RBPs (Baltz et al., 2012).

The RNA regulatory modules exhibited different patterns of RNA processing, degradation, localization, and translation. We speculate that these differences in RNA metabolism were driven by individual RBPs or the combination of RBPs associated with that regulatory module. This was supported by the response of specific RNA regulatory modules to ELAVL1 knockdown (Figure 4A, B). Additionally, the RNA regulatory modules encoded functionally related proteins and similarly localized proteins. The enrichments were for proteins with similar molecular functions or multi-component complexes rather than signaling pathways (Supplemental Fig 3b). Altogether, these lines of evidence provide support for the coordinate regulation of ‘functionally coherent’ RNA regulatory modules as proposed by the post-transcriptional operon/regulon model (Keene, 2007). The ultimate test of this model would involve manipulating specific combinations of binding sites and RBPs. Our study provides the rationale for such experiments, which unfortunately remain technically challenging.

Our observations have important implications for RBP-RNA regulatory networks and their importance in gene expression. The mRNA targets within specific regulatory modules encoded the RBP themselves, a generalization of a commonly made observation that RBPs bind to the mRNAs encoding them (Mesarovic et al., 2004). Our analysis lends support for this frequently observed potential auto-regulatory feedback. These feedback loops may in fact buffer the expression range of the targeted mRNAs, including those of the RBP. In this context, the observation that the RNA regulatory modules exhibited lower cell-to-cell gene expression variance, provides more evidence for the importance of post-transcriptional regulation in buffering transcriptional noise (Bahar Halpern et al., 2015; Battich et al., 2015). Systematic perturbation of individual and combinations of RBPs will be quite powerful in revealing fundamental properties of RNA regulatory networks such as auto-regulatory feedback and buffering.

The binding preference and targets of the vast majority of human RBPs remains unknown. The insights gained from this study demonstrate the value of large-scale efforts by ENCODE and others in the community to globally identify RBP binding sites. Of the 64 RBPs in this study, 44 were not represented in the ENCODE cell lines. Cumulatively these efforts interrogate ~10% of human RBPs with known RNA-binding domains. Thus, these two large scale efforts offer the potential to complement one another in our continuing attempts to understanding RBP-RNA regulatory networks, for which we have only glimpsed the tip of the iceberg.

## Acknowledgements

U.O. and T.T. acknowledge support from an award from the US National Institutes of Health (R01-GM104962). M.S. was supported by the ETIUDA scholarship # 2014/12/T/NZ1/00497 from National Science Center, Poland. N.M. acknowledges support from EU Marie Curie IIF, and the RNA Bioscience Initiative for startup funds.

## Author Contributions

Conceptualization, N.M., T.T, and U.O.; Methodology, N.M., H.W, M.S., M.G, A.G, A.M., T.T, U.O; Investigation, N.M., H.W, M.G, A.G, A.M., T.F, J.I.H, K.A; Formal Analysis, N.M., H.W, M.S., M.G, A.G, A.M.; Writing – Original Draft, N.M., T.T, and U.O.; Writing – Review & Editing, N.M., S.V. A.G. M.S., T.T, and U.O.; Funding Acquisition, N.M., T.T, and U.O.; Resources, N.M., T.T, and U.O.Supervision, N.M., T.T, and U.O.

## STAR Methods

### Processing, filtering, and quality control of PAR-CLIP libraries

Each PAR-CLIP library was subject to two rounds of quality control. First, all PAR-CLIP libraries generated in HEK293 cells were subject to the quality control pipeline PAR-CLIP Suite v1.0 (https://rnaworld.rockefeller.edu/PARCLIP_suite/). Using raw Illumina sequencing data, this pipeline identified the predominant target RNA category or categories for each RBP and provided the T-to-C conversion frequency resolved by read length and RNA category (Supplemental Fig 1). The mapped reads of each RNA category were resolved by error distance 0 (d0), error distance 1 (d1; split in T-to-C and d1 other than T-to-C), and error distance 2 (d2). This process discriminated for each library true target RNA categories from non-crosslinked background RNA categories populated by fragments of abundant cellular RNAs. In order to disqualify experiments comprising too many non-crosslinked RBP-specifically bound RNAs or co-purified non-crosslinked background RNAs, we pursued only datasets which collect at least 10,000 redundant d1 reads ≥ 20 nt in at least one of major RNA annotation categories with d1(T-to-C)/(d0 + d1) ≥ 30%, and d1(T-to-C)/(d1-total) ≥ 65%.

For the libraries passing the first threshold, we defined and annotated binding sites using PARpipe, which is a pipeline wrapper for PARalyzer (Corcoran et al., 2011; Mukherjee et al., 2014). The threshold for additional filtering were determined by comparisons with the reference library (Friedersdorf and Keene, 2014). This reference library was generated using a modified PAR-CLIP protocol in which there was no immunoprecipitation and the addition of an rRNA depletion step after proteinase K digestion, followed by a partial digestion using RNase T1. We required libraries had to have an average fraction T-to-C over remaining reads greater than 0.32 (the average fraction T-to-C over remaining reads greater of the reference library), an average conversion specificity greater than 0, more than 20000 aligned reads, not be digested only with micrococcal nuclease, a redundant read copy fraction less than .98 (Supplemental Fig 1b,c and Sup Table 1). For RBPs with three or more libraries, we removed outlier based on correlation of 6-mer frequency calculated from PARalyzer-utilized reads.

### Annotation category preference and positional analysis of binding density

For calculating the annotation category preference, we calculated the difference in the fraction of T-to-C reads per annotation category between each RBP library and the reference library. For example, if the fraction of miRNA annotated reads with T-to-C transitions in a specific RBP library was 0.20 compared to 0.05 in the reference library, the miRNA preference value for this specific RBP is 0.15. For the positional binding analysis, we selected genes (n=15120) using GENCODE v19 as annotation based on our earlier work on HEK293 RNA processing and turnover dynamics (Mukherjee et al., 2017). Isoform expression was calculated using RSEM (Li and Dewey, 2011). For each gene, we selected the transcript isoform with the highest isoform percentage or chose one randomly in case of ties (n=8298). The list of selected transcript isoforms was used to calculate the median 5’ UTR, CDS and 3’ UTR length proportions (5’ UTR=0.06, CDS=0.53, 3’ UTR=0.41) using R Bioconductor packages GenomicFeatures and GenomicRanges. For regions downstream annotated transcription ends (TES) and adjacent to splice sites, we chose windows of fixed sizes (TES 500nt, 5’ and 3’ splice sites 250nt each). We generated coverage tracks from the PARalyzer output alignment files and intersected those with the filtered transcripts. Each annotation category was binned according to its relative coverage averaged according to each bin. For intronic coverage, we averaged across all introns per gene, given a minimal intron length of 500nt. All bins were stitched to one continuous track per transcript. Altogether 6632 intron containing transcripts showed coverage in at least one PARCLIP library. For each library, we required transcripts to have a minimal coverage maximum of > 2. For each transcript, we scaled the binned coverage dividing by its maximal coverage (min-to-1 scaling) to emphasize spatial patterns independent from transcript expression levels. Replicate RBP PARCLIP libraries were combined at this point. Transcripts targeted in more than one replicate library were aggregated using the average of their binned coverage. RBPs with less than 50 filtered target transcripts (after aggregation) were not considered. Next, we split transcript coverage in two parts, separating 5’ UTR to TES regions and intronic regions. To generate the scaled meta coverage across all targeted transcripts per RBP, we used the heatMeta function from the Genomation package. For the 5’UTR to TES, we scaled each RBP meta-coverage track independent of other RBPs. For each RBP, we subtracted the scaled meta coverage of PARCLIP reference library (Friedersdorf and Keene, 2014). For intronic sequences, we scaled each RBP relative to all other RBPs to highlight RBPs with more substantial intronic binding patterns. Finally, we visualized the density using pheatmap.

### Sequence analysis

We calculated 6-mer frequencies with Jellyfish from all reads that generated a PARalyzer binding site for each library. For each RBP, we selected the library with the lowest percent of duplicated sequences (see supplemental table 1) to serve as a representative library for the sequence analysis and factor analysis. For each RBP, we counted the number of 6-mers with a frequency of x or higher, where x was from 1/4096 to 1/4. To evaluate the 6-mers enriched by a given RBP relative to the reference library, we regressed the RBP 6-mer frequency against the the reference library 6-mer frequency and collected the residuals (the unexplained variance). Next, identified all 6-mers that were found as the top 5 enriched over the reference library for any of the analyzed RBPs. We clustered the enrichment scores for the 6-mers across all RBPs and generated a heatmap using the ‘aheatmap’ function in NMF R package. We ran SSMART using all binding sites found in mRNA-derived annotation categories ranked by the library size normalized enrichment over the reference library.

### Factor analysis

For each site identified we calculated a library size normalized enrichment compared to the the reference library. We calculated the sum of all enrichment scores for all sites annotated as mRNA and lncRNA. Next, we normalized for expression levels (collected the residuals) to create the final matrix of values. The number of factors, 10, was determined using the majority result of numerous methods to estimate the number of factors. Clustering of the score matrix was performed using the most stable results from numerous iterations of k-means clustering.

### Gene ontology analysis

Multiple-test corrected gene ontology enrichment values were calculated using the TOPGO R package. For each set of genes, we used all 13,299 genes in the factor analysis as the background or gene universe. Enrichment was calculated using the ‘parent-child’ approach on the top 100 enriched terms. This metric accounts for the hierarchical organization of gene ontology terms to minimize false-positive enrichments. We performed a Bonferonni multiple test correction on the enrichment p-values.

### Premature and mature RNA quantification

Mature- and premature-transcript expression, transcripts per million (TPM), was quantified with RSEMv1.2.11 (http://deweylab.biostat.wisc.edu/rsem/src/rsem-1.2.11.tar.gz) as described previously (Mukherjee et al., 2017). Briefly, for each gene we included an additional isoform corresponding to the sequence of the full gene locus. Specifically, we modified the GENCODEv19 gtf and used this as the input for the ‘rsem-prepare-reference’ function to generate a modified index used for quantification. For each gene, we calculated the expression of ‘mature’ RNA as the sum of all isoforms for that gene excluding the ‘primary’ transcript. For intronless genes, premature and mature expression values were summed. We performed this analysis on the ELAVL1 knockdown RNA-seq experiments (Kishore et al., 2011).

### Cell-to-cell expression variability

RNA-seq gene expression data for 25 individual HEK293 cells were downloaded from (Guillaumet-Adkins et al., 2017). We calculated the coefficient of variation (100*standard deviation/mean) for each gene across all 25 cells.

---

**Figure S1:**
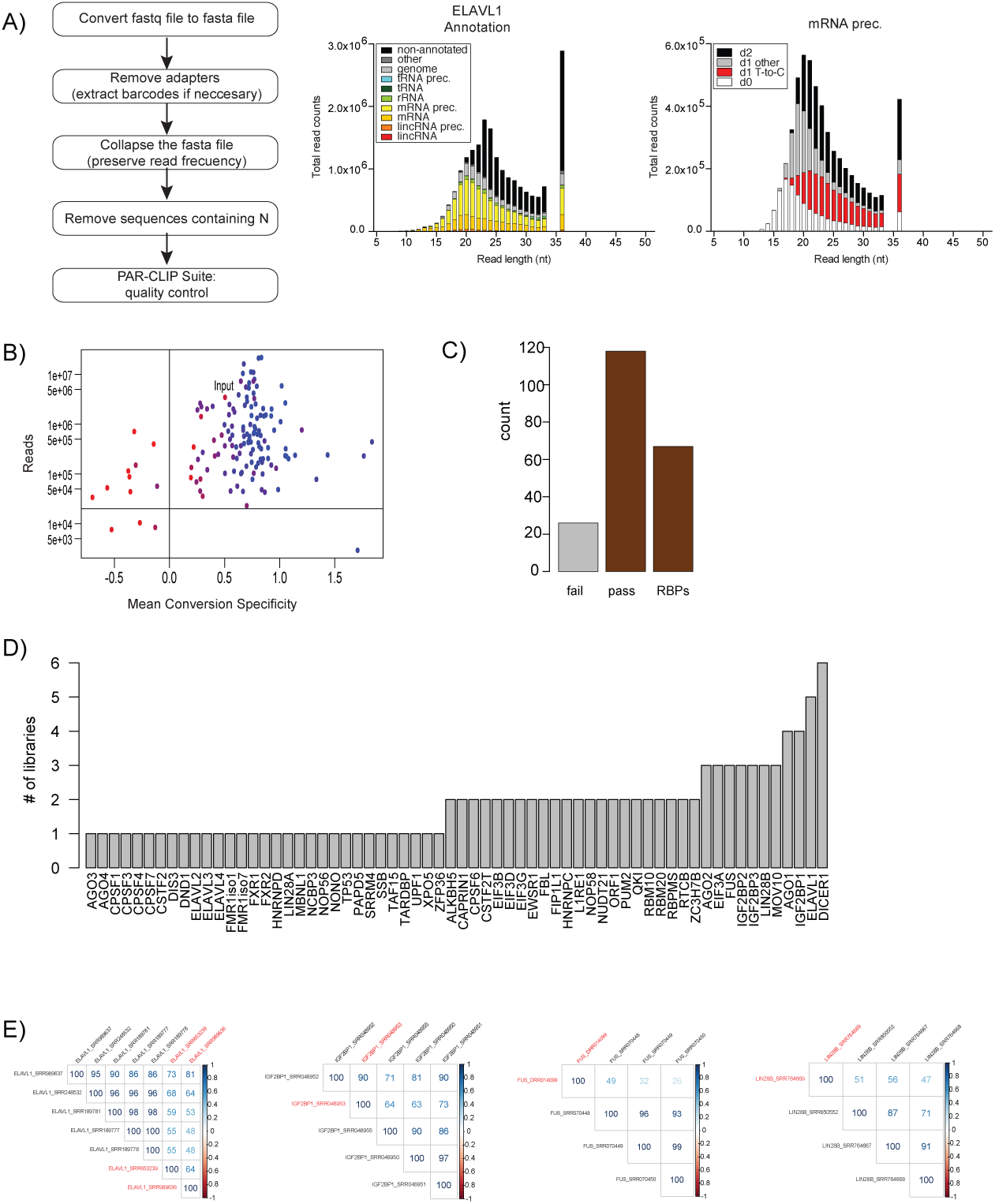
QC filtering of libraries. A) Description of PAR-CLIP suite to assess library quality control per annotation category (left). Example of number of reads mapping to each RNA category with up to 2 mismatches resolved by length of adapter-extracted sequence reads for an ELAVL1 library (middle). Sequencing read composition of the most abundant RNA category fir the ELAVL1 library. Reads were assigned as d0 (white), d1 T-to-C (red), d1 other than T-to-C, (light gray), and d2 (black) (right). B) Libraries had to have > 20,000 aligned reads and a mean conversion specificity > 0, and a higher mean T-to-C fraction than the the reference library (red lower, blue higher). C) Number of libraries analyzed and their quality control status. D) Count of libraries passing QC per RBP. E) Examples of outlier library removal (libraries labeled with red text were removed) based on correlation of read 6-mer frequency for RBPs with 3 or more libraries.

**Figure S2:**
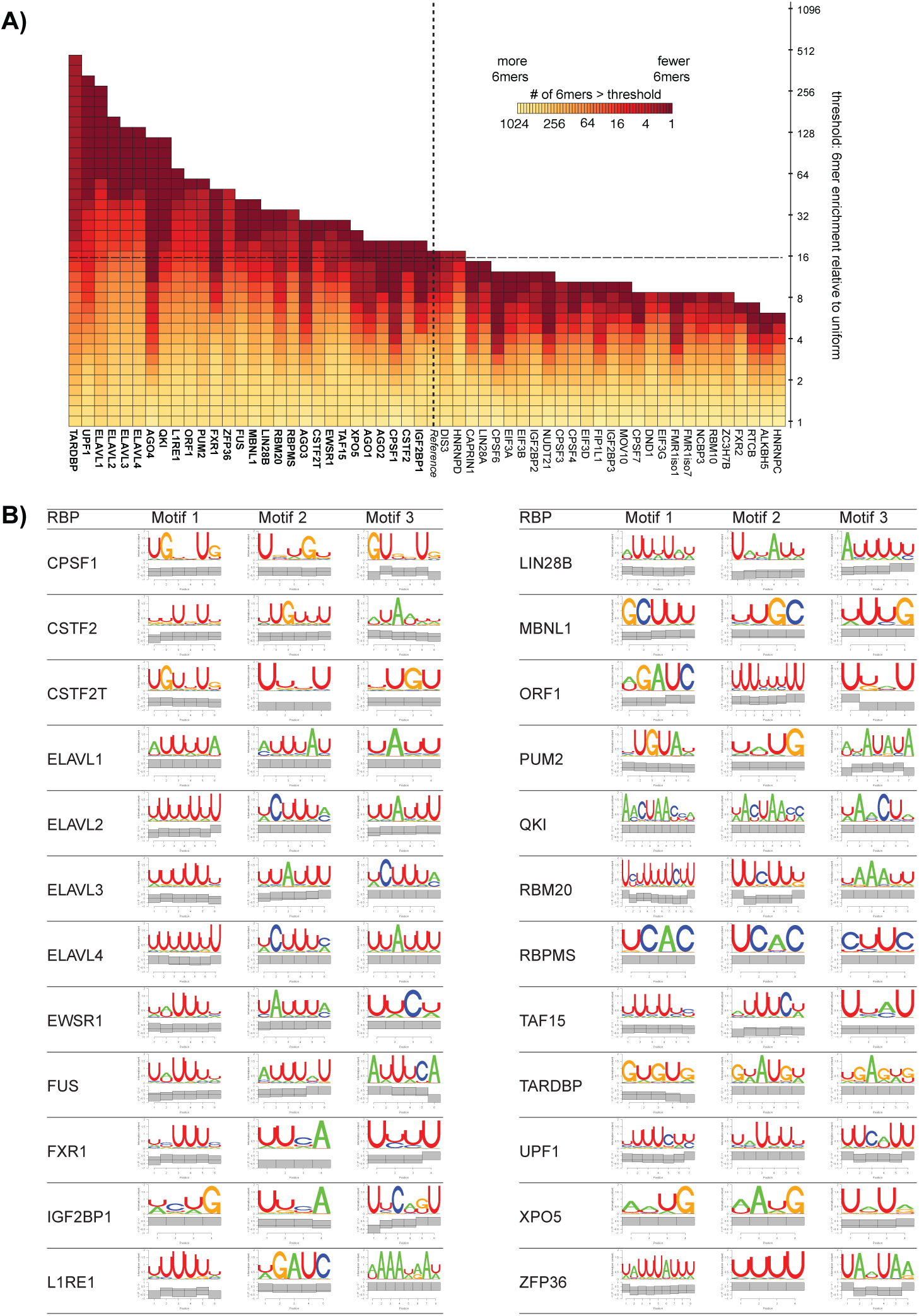
Grouping RBPs by sequence specificity. A) Heatmap of the number of 6-mers enriched per RBP at different specificity thresholds. The color scale represents the log_2_ [number of 6-mers] that are enriched at a given threshold (y-axis). The thresholds are represented as log_2_ [6-mer frequency]. There are 4096 different 6-mers and if they were uniformly present this would represent a value of - 12 =log_2_ [1/4096]. The horizontal dashed lines at -8, represents 16-fold enrichment over a uniform background. For reference, the vertical dashed lines indicate the behavior of the reference library. B) Top 3 SSMART motif results using all binding sites found in mRNA-derived annotation categories ranked by the library size normalized enrichment over reference library.

**Figure S3:**
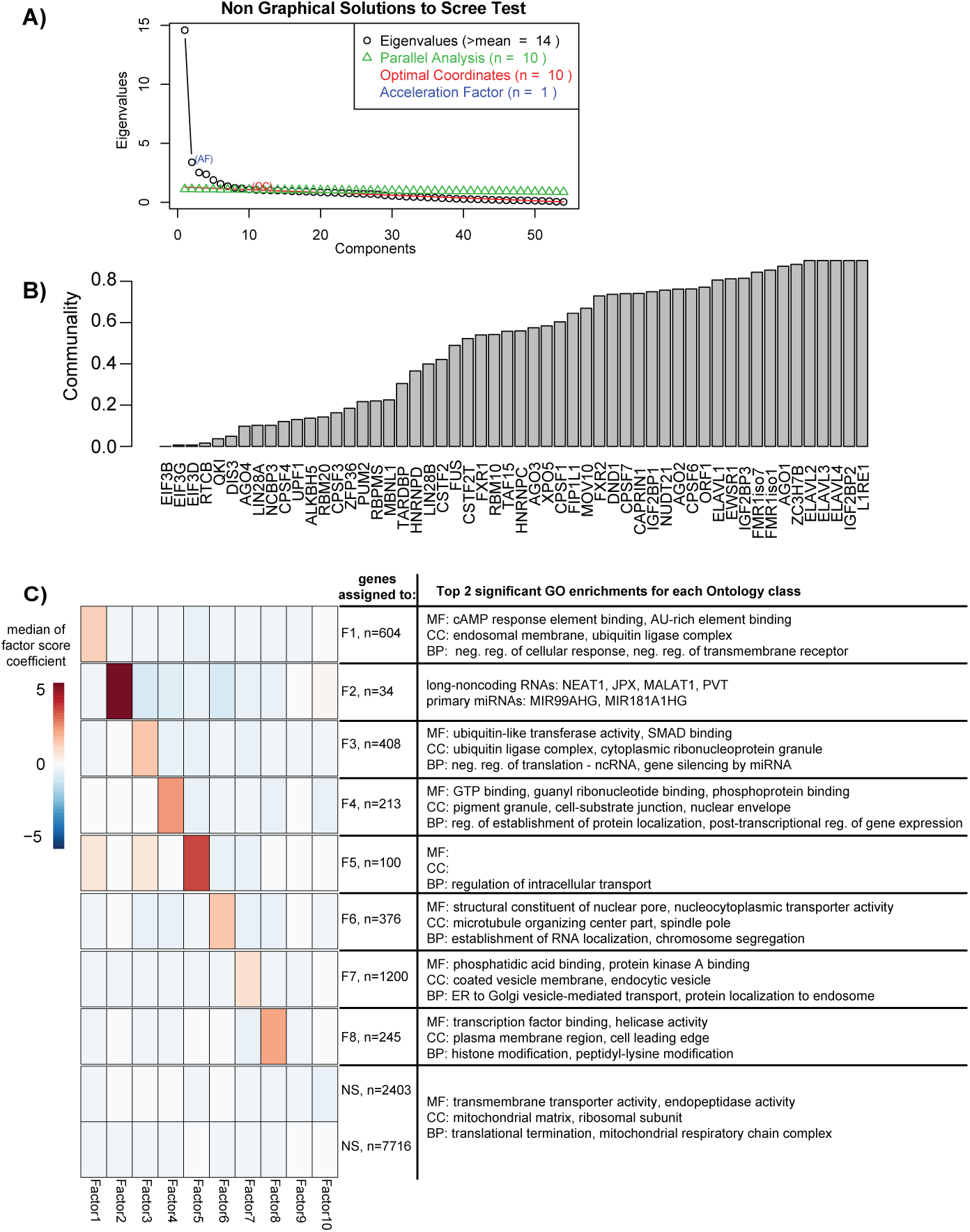
Factor analysis model selection and performance. A) Plot of eigenvalues versus number of factors to determine the optimal number of factors using four methods (different colors). B) Barplot of the communality, or the variance in a given RBP cumulatively explained by the all factors. C) Heatmap of the median factor score coefficient value for all genes that clustered together. The number of genes assigned to a specific factor and the top two most significant enriched GO annotations for each ontology class: molecular function (MF), cellular component (CC), and biological process (BP).

**Figure S4:**
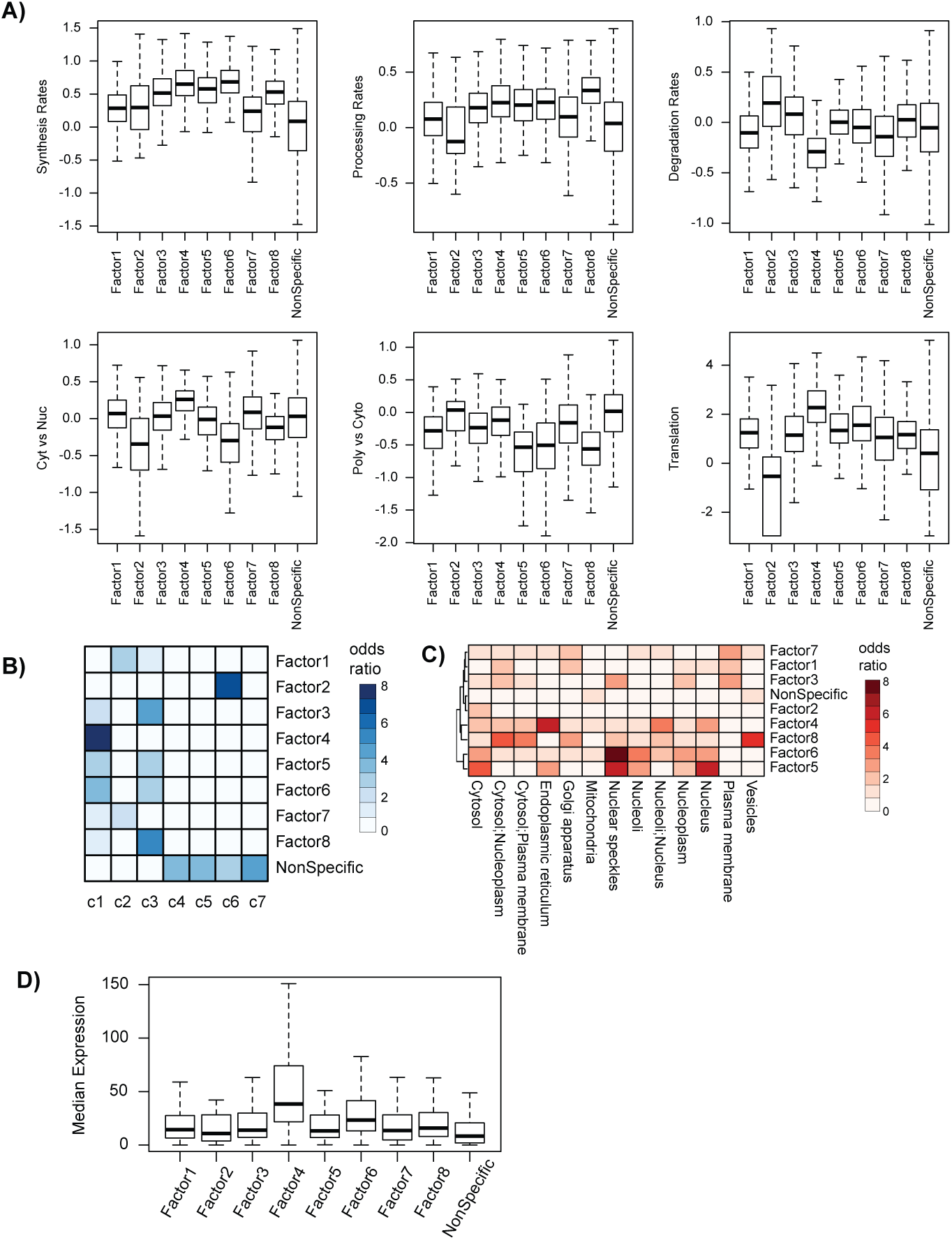
RNA metabolism profiles for factor-associated gene sets. A) Box-and-whisker plot for each gene set of the synthesis rates, processing rates, degradation rates, cytoplasmic versus nuclear localization (Cyt vs Nuc), polyribosomal versus cytoplasmic localization (Poly vs Cyt), and translational status from ribosome profiling data. B) Heatmap of the odds-ratio of the overlap between factor associated gene sets with RNA categories based on similar metabolic profiles from (Mukherjee et al., 2017). C) Heatmap of the odds-ratio of the overlap between factor associated gene sets and protein localization annotation. D) Box-and-whisker plot for each gene set of the median expression across 25 HEK293 cells.

